# Small Molecules Attenuate the Interplay between Conformational Fluctuations, Early Oligomerization and Amyloidosis of Alpha Synuclein

**DOI:** 10.1101/177006

**Authors:** Amrita Kundu, Sumanta Ghosh, Krishnananda Chattopadhyay

## Abstract

Aggregation of alpha synuclein has strong implications in Parkinson’s disease. The heterogeneity of folding/aggregation landscape and transient nature of the early intermediates result in difficulty in developing a successful therapeutic intervention. Here we used fluorescence measurements at ensemble and single molecule resolution to study how the late and early events of alpha synuclein aggregation modulate each other. The in-vitro aggregation data was complemented using measurements inside live neuroblastoma cells by employing a small molecule labeling technique. An inhibitor molecule (arginine), which delayed the late event of amyloidosis, was found to bind to the protein, shifting the early conformational fluctuations towards a compact state. In contrast, a facilitator of late aggregation (glutamate), was found to be excluded from the protein surface. The presence of glutamate was found to speed up the oligomer formation at the early stage. We found that the effects of the inhibitor and facilitator were additive and as a result they maintained a ratio at which they cancelled each other’s influence on different stages of alpha synuclein aggregation.

## INTRODUCTION

Protein aggregation is believed to be a common molecular theme for several neurodegenerative diseases, including Parkinson’s disease (PD), in which alpha synuclein (α-syn) aggregates to form Lewy bodies [1]. There is no cure available at present, and the heterogeneity of the folding/aggregation landscape is a crucial bottleneck towards the development of a disease-modifying therapeutics. Literature results suggest that the folding/aggregation landscape of α-syn is complex, containing at least three components. The most widely studied one is the formation of amyloid fibrils, which happens late. Amyloid fibril formation of α-syn and other proteins can be studied conveniently using thioflavin T (ThT), a dye whose fluorescence enhances when bound to the fibrils. While the late fibril formation can be investigated easily, monitoring early events is difficult. Using single molecule spectroscopy, it has been shown that the protein fluctuates rapidly between different conformers and form small molecules oligomers. We emphasize that it is important to study both the early and late events of aggregation. This is because; a) the early events may enable the faster detection of the disease and b) recent results point towards the toxicity associated with the early oligomers [2]. It is often argued that the formation of the amyloid fibrils may be a preventive mechanism by which toxicity induced by the oligomers could be minimized. However, opposing views exist, hypothesizing a direct role of amyloid fibrils (and Lewy Bodies) in PD pathology [3].

In this paper, we used a simple small molecules based strategy in combination with fluorescence spectroscopy at the ensemble and single molecule resolution to investigate both the early and late stages of α-syn aggregation. The objective of this study is to determine how the early and late stages of α-syn aggregation modulate the effect of each other. The first step of this strategy was to select two small molecules, which would either delay (the inhibitor) or promote (the facilitator) the overall aggregation of α-syn (the late stage outcome) in aqueous solution and inside live cells. The second step of the strategy was to probe the effects of the inhibitor and facilitator molecules on the early stages of α-syn folding/aggregation landscape. By doing these experiments, we aimed to find out if the early conformational fluctuations and oligomerization were affected by the inhibitor and facilitator molecules.

For the measurements of the late stage aggregation (the first step of the strategy), we used ThT fluorescence, which was complimented by dynamic light scattering (DLS) and atomic force microscopy (AFM). For the measurements inside live cells, we employed a cell culture model using small molecule fluorescent labels of α-syn expressing in neuroblastoma cell lines. For the measurements of conformational fluctuations and oligomerization at the early stage (the second step of the present strategy), we used the help of fluorescence correlation spectroscopy (FCS), a method which monitors diffusional and conformational dynamics of fluorescently labeled biomolecules at single molecule resolution. For the analysis of FCS data, the conventional discrete component analysis was supplemented by maximum entropy method (MEM).

We assumed that the late stage experiments in aqueous solution would enable us to determine the direct role these small molecules play to modulate the aggregation of the protein. These direct roles may originate from the binding between the protein and small molecule or from the preferential exclusion. In contrast, live cells data are expected to be complex containing multiple components of differing contributions. Protein aggregation, neuro-degeneration and the role of reactive oxygen species (ROS) have been extensively studied and widely debated. We used two amino acids, namely glutamate, and arginine as our choice for the facilitator and inhibitor respectively for the late stage of α-syn aggregation. These molecules are chosen for their proven roles in neurodegeneration. Glutamate, being an excitatory amino acid, has been shown to induce neurotoxic effects and time-dependent cell deaths [4-6]. Extracellular glutamate binds to its specific receptors resulting in calcium influx inside cells, which leads to the generation of ROS [4]. Interestingly, the accumulation of ROS has been shown to result in aggregation. It has been suggested by Hashimoto and Souza group that oxidative stress increases the accumulation and aggregation of α-syn inside cell [5, 6]. Long-term exposure to oxidative stress has been shown to induce α-syn aggregation in primary neuronal cultured cells [7]. Increased α-syn aggregation after rotenone exposure in cultured cells might be the result of increased ROS generation in the presence of rotenone. Although glutamate-induced neurodegeneration has been investigated, its effect on aggregation has not been studied in live cells. In contrast, L-arginine has been found to offer potent in-vitro and in-vivo neuroprotection [8]. Poly-arginine and arginine rich peptides have been shown to induce protection against glutamate induced excitotoxicity[9]. While the double role of amino acids (neurodegeneration vs. protection) has been a subject of extensive interests, any mechanistic understanding is absent.

The results obtained in the present study showed that the competition between conformational fluctuations and oligomerization could be a key factor in defining the folding-aggregation landscape of α-syn. The presence of arginine and glutamate influenced the competition between conformational fluctuations and oligomerizations differently. While the presence of arginine favored the compact conformer in equilibrium, the addition of glutamate sped up oligomers formation. We determined that the influence of one of these molecules can be canceled by introducing the second molecules. Finally using isothermal calorimetry (ITC) and mass spectroscopy, we found out that the effect of arginine was probably due to the binding of multimeric arginine onto the protein molecules, while glutamate remained excluded.

## RESULTS AND DISCUSSION

### *In-vitro* and live cell data suggest that arginine is an inhibitor and glutamate is a facilitator for the late stage of aggregation

First, we wanted to investigate how glutamate (which typically induces neurodegeneration) and arginine (often used as a neuro-protector) influenced the aggregation of α-syn. Since the measurement of the late stage of aggregation is relatively straightforward, it was attempted first. To induce aggregation, we agitated protein samples at 37°C for multiple hours, while the extent of aggregation was studied using ThT fluorescence. Normal glutamate concentration in blood is 40-50μM, and the average daily intake of glutamate in human body has been estimated to be 0.3-1 g [10]. That means that an average person of 70 kg can uptake around 0.06-0.2 mg/ml or 0.3-1.1mM of glutamate in blood, which is often utilized in glutamate-glutamine cycle in presynaptic terminals and glial cells[11]. In contrast, the use of arginine for the biological/clinical investigations has been well established, with the recommended dosage varying between μM and mM[12]. We studied the effect of the chosen amino acids at a wide range of concentration, which varied between 500μM and 200mM. Figure 1A,B,C show the variations of normalized ThT fluorescence intensity with time in the absence (black) and presence of different concentrations of arginine (blue) and glutamate (red). Figure 1D show the variation of t_lag_ in the presence of different concentrations of glutamate (red) and arginine (black). Since the parameter t_lag_ is an insensitive parameter, which was determined by mathematical fitting of the data (equation 1), we needed to use relatively high concentration of the amino acids in order to obtain a significant difference. We found that t_lag_(equation 2) decreased significantly in the presence of glutamate. In contrast, the presence of arginine increased t_lag_.

**Figure 1:**
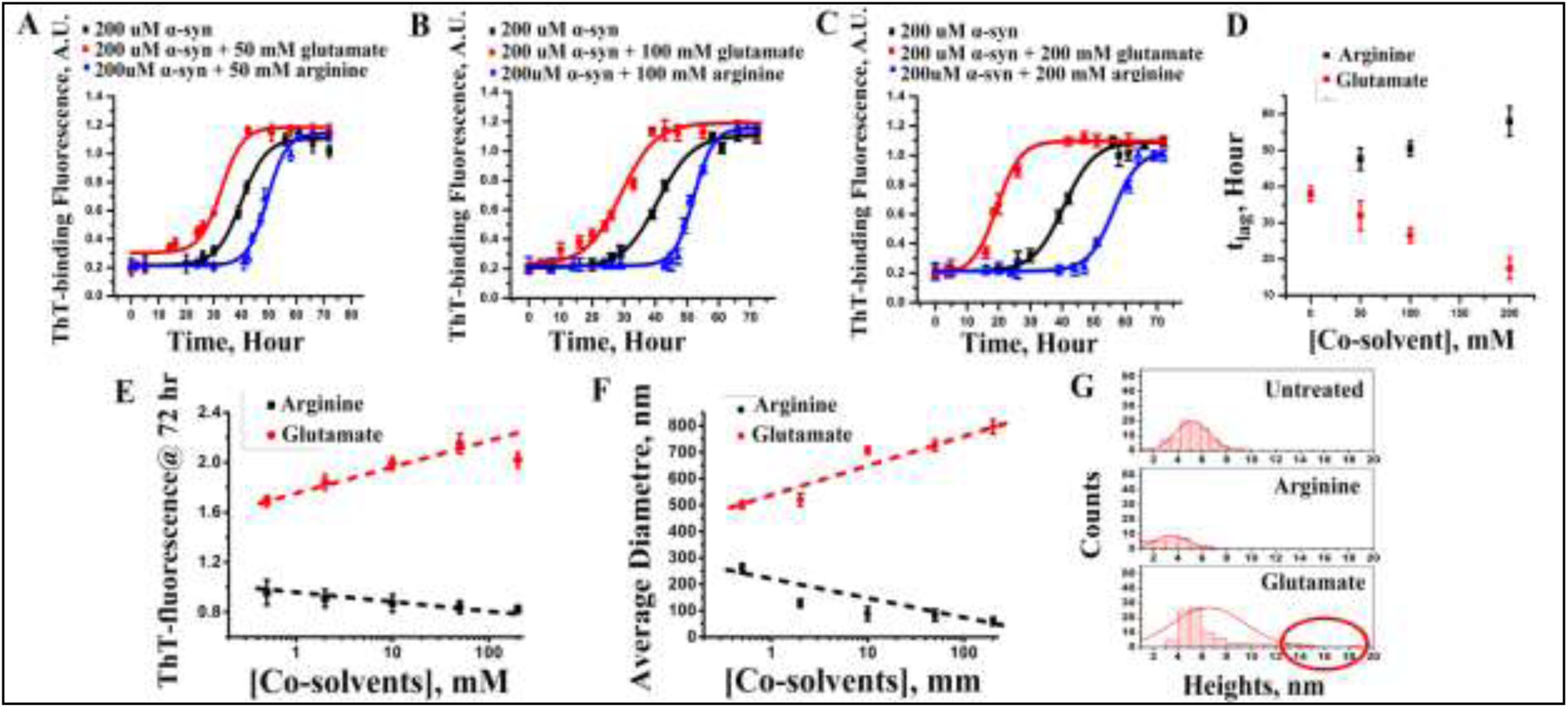
[A] ThT-binding fluorescence with 50mM, [B] 100mM and [C] 200mM arginine(blue) and glutamate(red) concentrations. ThT-binding fluorescence of wild type α-syn had been shown as a control (black) set of data in these figures; Data shown are mean±standard error. Error bars of ThT fluorescence experiments are calculated after repeating the experiments independently for eight times. [D] The variations of t_lag_of amyloidosis with arginine and glutamate concentrations are also shown. [E] Plot of ThT fluorescence at the saturating condition (72 hour)with increasing concentrations of arginine(black) and glutamate(red) from 500μM to 200mM.[F] Plot of average diameter in nm (from DLS) at the saturating condition with increasing concentrations of arginine(black) and glutamate(red) from 50μM to 200mM. For both E and F, x-axes were plotted using log scale, and hence the values at zero concentration are not shown, [G] Histogram of counts and height of α-syn fibrils at the saturated phase calculated from AFM data. Circle shows appearance of fibrils with heights ranging from 12-20nm.

The non-normalized ThT fluorescence kinetics are shown in Figure S1A-C. We plotted non-normalized ThT fluorescence intensity data obtained at the saturating incubation condition (72 hours) in Figure 1E, which showed significant concentration dependence between 500μM and 200mM. Figure 1E suggests that the extent of fibril formation was greater in the presence of glutamate, while the presence of arginine decreased the extent of fibrillation. Since ThT may not bind any non-fibrillar aggregate forming at the late stage, the extent of aggregation at the saturating condition (72 hours) was further studied using dynamic light scattering (DLS) and atomic force microscopy (AFM), which confirmed the above trends (Figure 1F, G). Detailed DLS and AFM data in the presence of glutamate and arginine are shown in FigureS1D-F.

We complemented above *in vitro* results in live cells expressing α-syn. For the studies in mammalian cells, we initially made a GFP-tagged α-syn construct and transiently transfected HeLa and SH-SY5Y cells, which did not show any fluorescent aggregate in either cell line (Figure 2A). Since the molecular weight of GFP (∼27kDa) is greater than that of α-syn (∼14kDa), the influence of GFP on the aggregation property of α-syn may not be negligible [13]. To circumvent this problem, we used a small ∼ 1.5kDa tetra-cysteine tagged α-syn (TC-AS), which can be labeled using FlAsH-EDT_2_, for the aggregation studies inside live cells [14, 15].A 12-mer peptide, with four cysteine residues and one proline and glycine in the middle (– CysCysProGlyCysCys-), has been fused with the C-terminal end of α-syn (TC-AS). This TC-AS containing pcDNA3 plasmid has been transfected to HeLa and SHSY-5Y cell lines. This small 1.5kDa tag is non-fluorescent but in the presence of FlAsH-EDT_2_, it turns into a fluorescent complex Detailed investigation on WT α-syn and TC-AS (in the absence and presence of fluorescence labels) suggested identical aggregation behavior for these systems [16]. We found that TC-AS started expressing in SH-SY5Y cells after 24 hours of transfection (Figure 2B). The formation of bright fluorescent aggregates became visible in the cytoplasmic area with 50-60% of the cells after 48 hours. After 72 hours of transfection, the presence of vacuoles was observed in aggregates containing cells.

**Figure 2:**
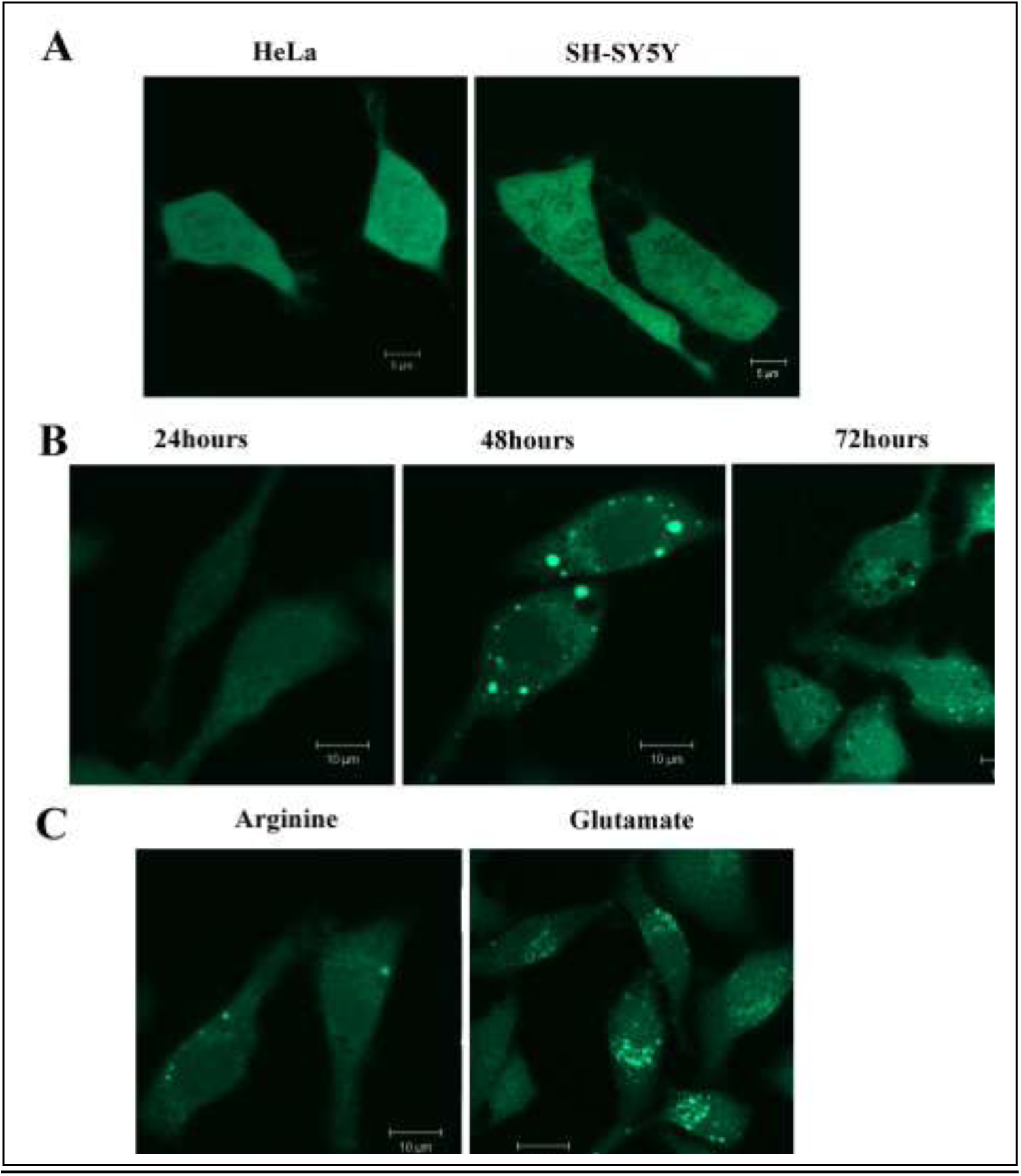
[A] Confocal images of HeLa and SH-SY5Y cells expressing GFP-tagged α-syn after 3 hours of incubation at 37°C. [B] SH-SY5Y cells expressing TC-AS; 24 hours, 48hours and 72 hours after transfection. [C] SH-SY5Y cells treated with 500μM of arginine and glutamate. The imaging was performed on n=250 TC-AS over-expressing cells, one of them is shown above as a representative. Scale bars, 10μm.

Subsequently, we studied the effect of 500μM arginine and glutamate (Figure 2C) on the TC-AS transfected cells. From the images, we calculated three parameters to semi-quantitatively estimate the extent of TC-AS aggregation. These parameters are i) the diameter of an individual aggregation spot, ii) the maximum intensity of an individual spot, and iii) the integrated area of an inclusion. These parameters determined at different conditions are shown in Table 1. In the untreated cells, the average diameters of the fluorescent spots were calculated to be between 3 and 7μm. The corresponding diameters in the presence of glutamate and arginine were found to be greater than 8μm and around 1-2 μm respectively. The intensity profiles for the aggregates inside SH-SY5Y cells are plotted in Figure 3. In the presence of arginine (Figure 3B), the intensity of individual spots decreased significantly. Interestingly, glutamate treatment (Figure 3C) did not change the maximum intensities of individual spots (compared to the untreated conditions, Figure 3A). In contrast, the presence of glutamate resulted in the merging of individual spots forming large inclusions (Figure 3C). Similar observations were found in HeLa cells in the treated and untreated conditions (Figure S2). We subsequently calculated the integrated area of the inclusions in untreated and treated cells (Table 1). Figure 3D and Eshow the distribution of aggregate sizes at different conditions for both HeLa and SH-SY5Y cells.

**Figure 3:**
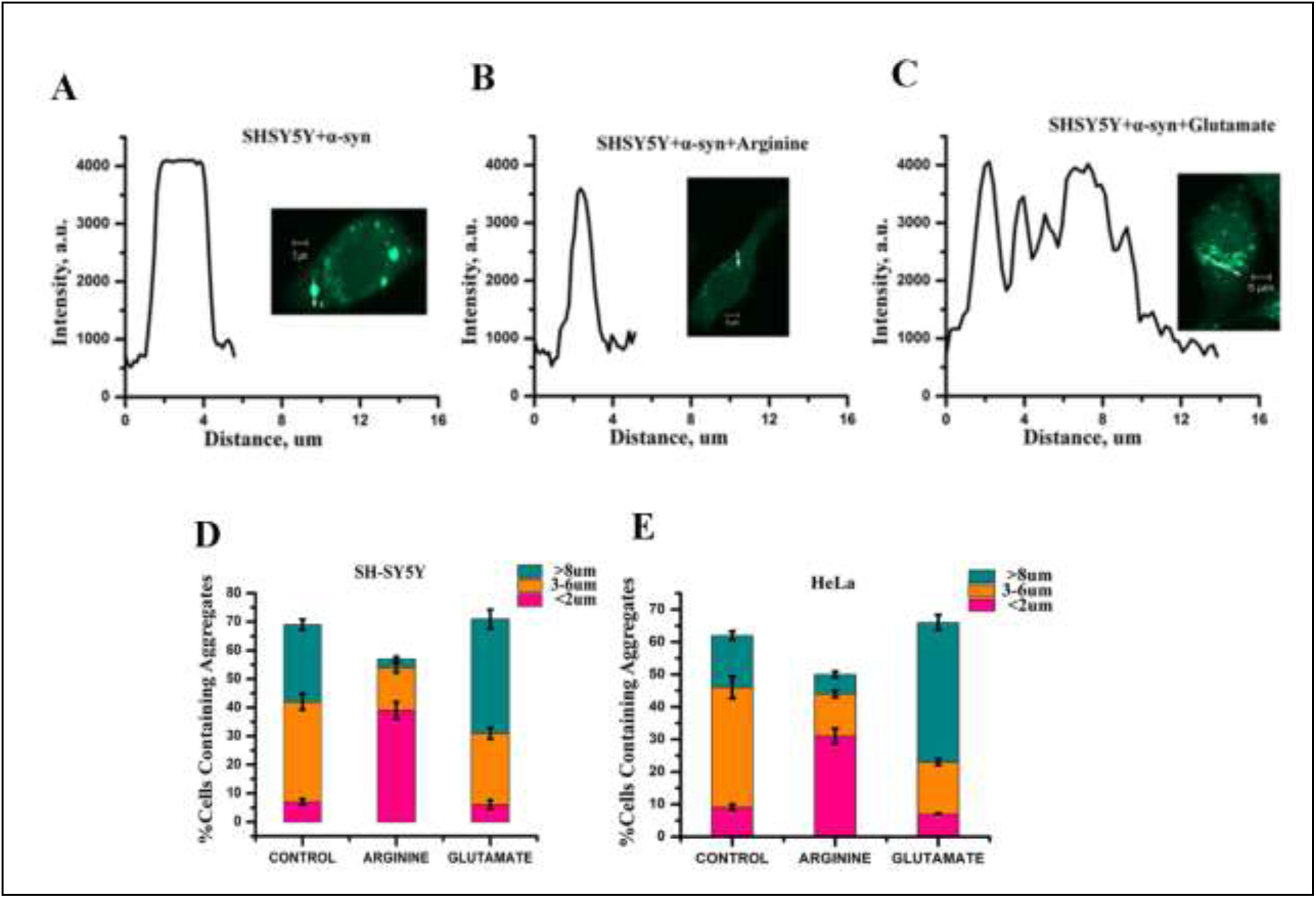
SH-SY5Y cells containing TC-AS aggregates [A] untreated, [B] in the presence of 500μM arginine, [C] in the presence of 500μM glutamate. The intensity values were calculated along the white arrow. These values as a function of distance are plotted at the left side of the images. These analyses were done for 10 individual cells for each condition and one of them is shown above as a representative. Scale bars, 5μm.The distribution of different sized TC-AS aggregation in [D] HeLa and [E] SH-SY5Y cells. Different colored regions represent the size of the aggregates; pink: aggregates smaller than 2 μM, orange: larger than 3μM but smaller than 7μM, cyan: larger than 8μM. Bars indicate average values of three independent experiments. Data shown are mean±standard error.

**Table 1:**
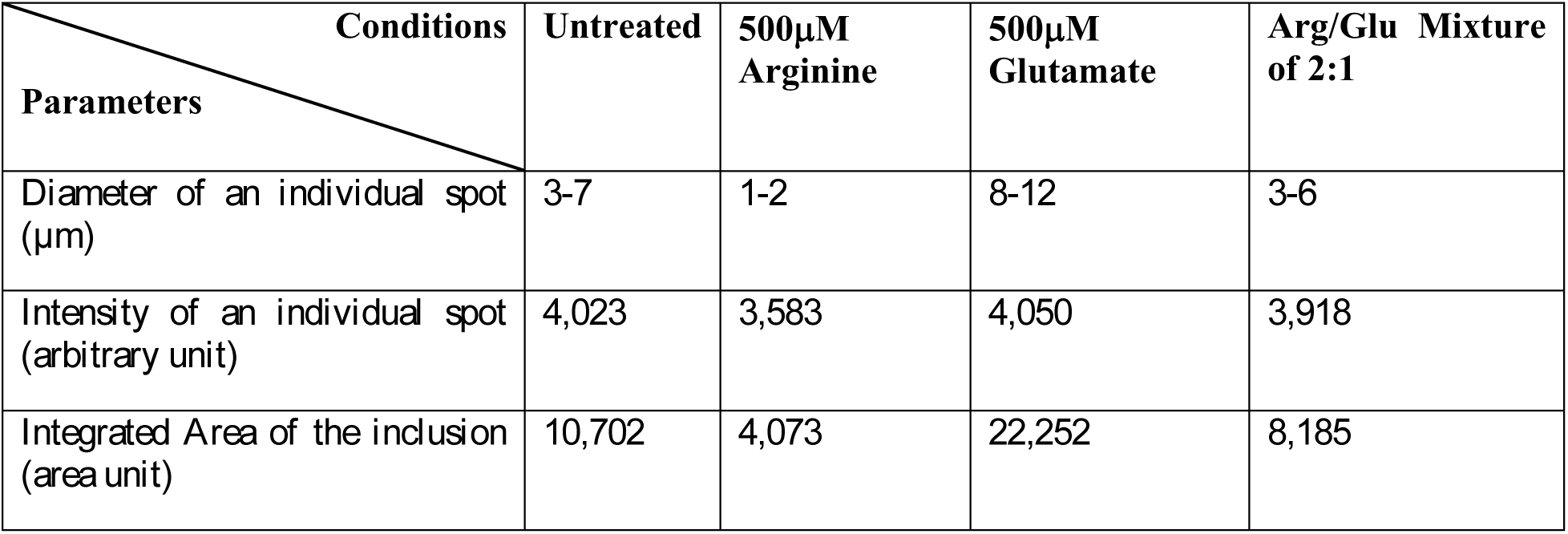
The effect of arginine and glutamate on TC-AS aggregation inside SH-SY5Y cell lines

Subsequently, we studied the effect of several other small molecules as controls to investigate their effects in vitro and inside cells. These included aspartate and lysine, which resembled glutamate and arginine respectively in terms of their electrostatics. Both in vitro and live cell measurements with lysine suggested significant increase in aggregation (although arginine showed the reverse trend) (Figure 4A-B). While aspartate is identical to glutamate in its charge characteristics, its effect of the former was significantly less both in aqueous solution (Figure 4A) and in cellular environment (Figure 4B).NaCl was also used as another control, as both arginine and lysine were used as their hydrochloride salts. We wanted to determine if the inhibition observed by arginine originated from the chloride ions present in solution. The results in aqueous solution showed that the addition of salt increased aggregation with increasing concentration of NaCl, with very high extent of aggregation observed in the presence of 100mM concentration. Since arginine contains a guanidinium moiety in its structure, which is expected to play an important role in the aggregation inhibition property (see later), we used guanidinium hydrochloride as another control. The presence of guanidinium hydrochloride resulted in inhibition of the aggregation, which was observed for both in-vitro and in cell experiments (Figure 4A and B). However, there was a significant difference in the behavior of arginine and guanidinium hydrochloride. While arginine data showed that the extent of aggregation decreased in a dose-dependent manner, the trend was reverse for guanidinium hydrochloride.

**Figure 4:**
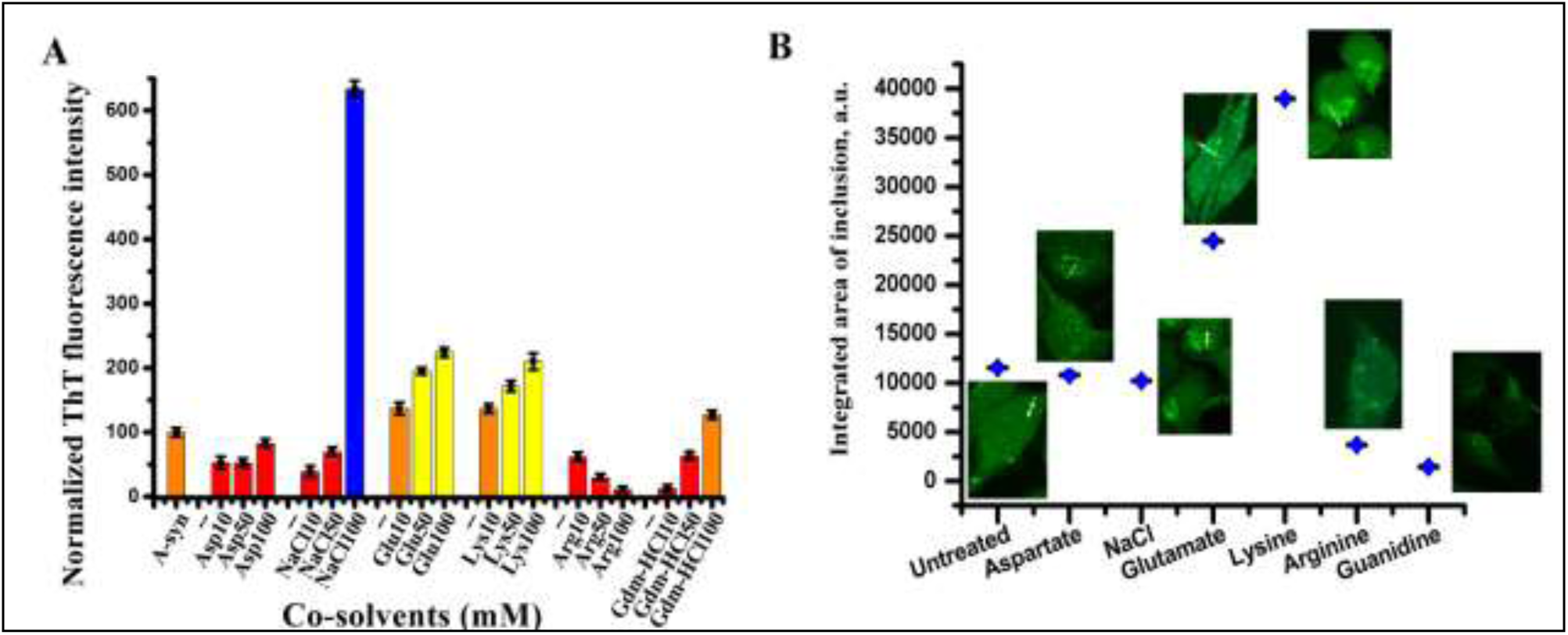
Aggregation screening of α-syn using different small molecules. [A] Normalized ThT fluorescence data of α-syn in the absence and presence of small molecules at different concentrations (10, 50 and 100mM). [B] The integrated area of α-syn inclusions inside SH-SY5Y cells have been plotted in the absence and presence of different small molecules at 50mM concentration.

We used fluorescence recovery after photo-bleaching (FRAP) to determine the mobile fraction (F_M_) and the diffusion coefficient (D) (equation 3) of α-syn inside cells (Table 2). As control, we chose a region, where inclusion free diffused expression of TC-AS was observed (Figure S3A). Interestingly even for the aggregate free regions, the fluorescent signal did not reach its initial value after bleaching (F_M_=51%) (Figure S3A, Table 2). This observation suggested relatively slow intercellular dynamics of the fluorescent TC-AS molecules (even when they were not aggregated). However, the recovery of the molecules at the aggregated regions (Figure S3B, Table 2) was considerably less (F_M_=19%) compared to that at the cytoplasmic regions (F_M_=51%). The recovery from the aggregate containing regions improved significantly in the presence of arginine (F_M_=35%) (Figure S3C, Table 2). In contrast, the presence of glutamate (Figure S3D, Table 2) resulted in decreased recovery (F_M_=14%).

**Table 2:** The values of mobile fraction and diffusion constant of TC-AS

| **Parameters \ Conditions** | **Percentage of Mobile fractions ($F_M$), %** | **Calculated Diffusion coefficient, μm<sup>2</sup>/s** |
| --- | --- | --- |
| Untreated cell cytoplasm | 51±4 | 0.077±0.00291 |
| Untreated cell inclusion | 19±3 | 0.049±0.00089 |
| Arginine treated cell inclusion | 35±5 | 0.061±0.0031 |
| Glutamate treated cell inclusion | 14±3 | 0.044±0.00127 |

A mono-exponential fit to the recovery phase of the FRAP data (equation 4) in the cytoplasmic aggregate free region yielded the D value of 0.077 μm^2^/sec. The value of D obtained in the present study is similar to the previously published results (0.08-0.15 μm^2^/sec [17]). The value of D at the aggregated region was 0.049μm^2^/sec, which was significantly less than the cytoplasmic D. The values of D of the aggregated molecules changed to 0.061μm^2^/sec and 0.044μm^2^/sec in the presence of arginine and glutamate respectively. Interestingly, the presence of arginine or glutamate in the inclusion free cytoplasmic area did not change their F_M_ or D values.

The viabilities of cells after incubated with the aggregated α-syn in the absence and presence of the co-solvents were measured by MTT assays[18]. Both SH-SY5Y and HeLa cells were incubated for 24 hours in the presence of α-syn aggregates at 10μM concentration. As shown in Figure S4A, the presence of aggregates increased cytotoxicity when compared to the normal cells. The cell viability decreased even further in the presence of glutamate. As expected, the addition of arginine showed improved viability. Nuclear counterstaining of α-syn aggregates containing SH-SY5Y cells with Hoechst showed round and intact nucleus indicating that the cells were in good health (Figure S4B).

### Arginine, the inhibitor, populates the compact conformer in early fluctuations, while glutamate, the facilitator, speeds up oligomer formation

In the previous section, we established that glutamate is a facilitator, which sped up the aggregation kinetics at the late stage. In contrast, arginine acted as an inhibitor of the late stage aggregation. We observed similar results *in vitro* and in two different cell lines. Subsequently, we studied the effects of arginine and glutamate on the early stages of α-syn folding/aggregation landscape. The idea was to investigate how the presence of facilitator glutamate (or inhibitor arginine), influenced the early events (conformational fluctuations and oligomerizations). For the measurements of early fluctuations and oligomerization and to separate out their contributions, we used FCS. A schematic diagram of a typical FCS setup is shown in Figure 5A. Any increase or decrease in molecular size (as a result of conformational change) would increase the hydrodynamic radius (r_H_) of the protein, keeping the number of particle (N) unchanged. In contrast, oligomerization would increase r_H_ while decreasing N.

**Figure 5:**
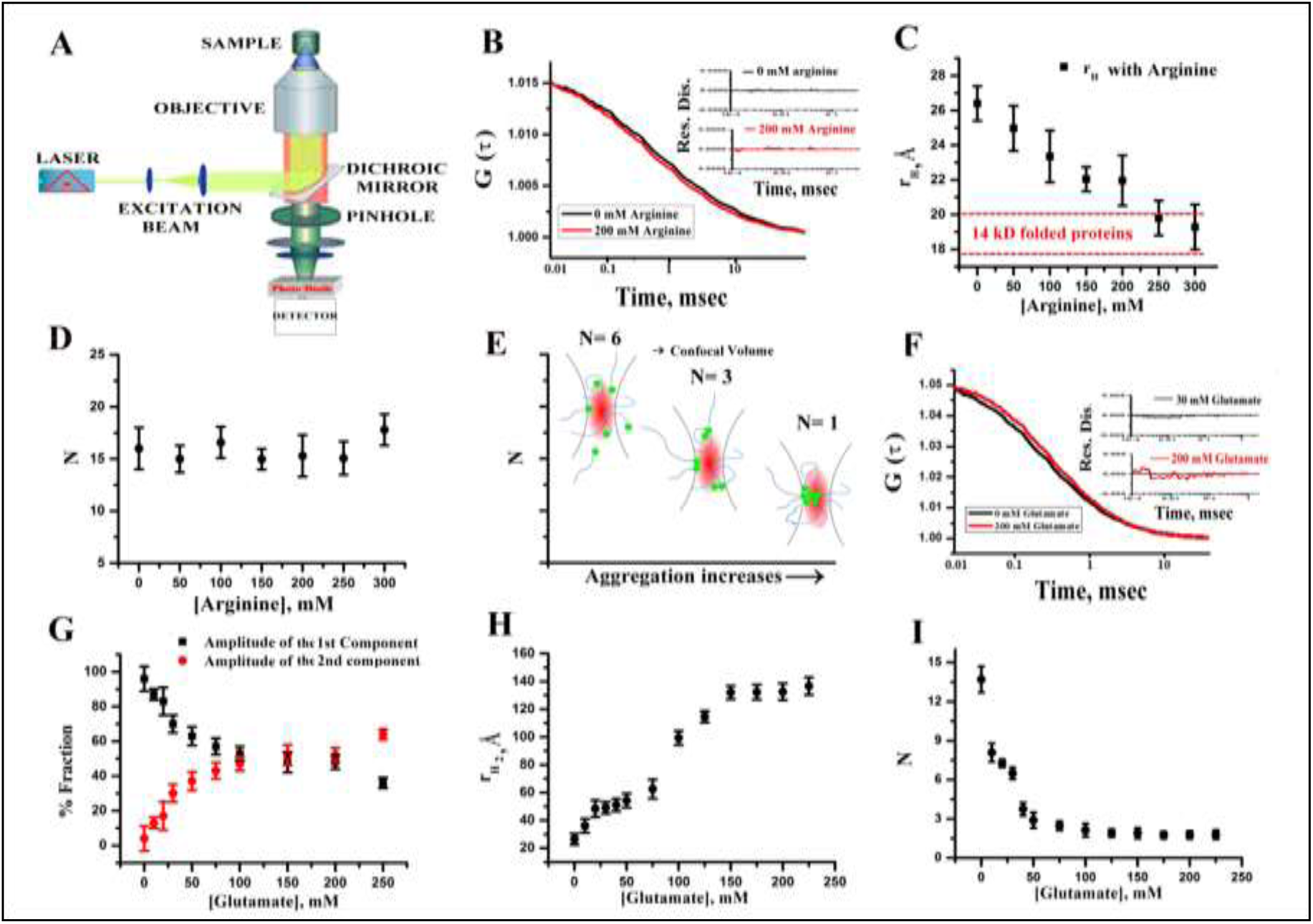
[A] The schematic diagram of a typical FCS setup [B] the variations of the correlation functions obtained in the absence (black) and presence of 200 mM arginine (red). Randomness of the residual distributions in the absence and presence of arginine are shown in the inset; [C] the variations in r_H_with increasing concentrations of arginine; [D] the changes in the average number of particles (N) with increasing concentrations of arginine; [E] a typical representation showing the decrease in N with aggregation; [F] representative correlation functions obtained with 0 and 200 mM of glutamate are shown here. The data were fit to a model assuming two components diffusion. Randomness in the residual distributions at different concentrations of glutamate are shown in the inset; [G] the variations of the amplitudes of two components as a function of glutamate concentrations; [H] the variations of r_H_with increasing concentrations of glutamate indicating the formation of oligomers; [I] the decrease in N as a function of increasing concentrations of glutamate also supports glutamate induced oligomerization; Data shown are mean±standard error. Error bars were calculated after five times independent repetition of the experiments.

For the present FCS experiments, we chose a single cysteine mutant of α-syn (G132C), which was labeled using Alexa488Maleimide (Alexa488). G132C α-syn is considered similar to wild type protein for biophysical studies[19, 20]. The Alexa488 labeled G132C protein would be referred as Alexa488Syn from now onwards. We studied diffusional dynamics of Alexa488Syn at pH 7.4 in the absence and presence of different concentrations of arginine and glutamate. These experiments were carried out under non-aggregating conditions (25°C, no incubation).

The correlation functions (representative data shown in Figure 5B) obtained with Alexa488Syn in the absence and presence of different concentrations of arginine, were fit using a single component diffusion model (equation7). The goodness of the fits was established using the randomness of the residual distributions (inset of Figure 5B). The value of r_H_(equation 11) was found to be more than that expected for a compact well-folded protein containing 140 amino acid residues (calculated to be 17.8-19.9 Å)[21], but less than that obtained with the unfolded state of similar proteins (calculated to be 33.3-36.9 Å). This observation was in accordance with the natively unfolded nature of the protein. With the addition of arginine, we saw a decrease in r_H_ indicating compaction of the natively unfolded structure (Figure 5C). The decrease in r_H_ (and of τD) in the presence of arginine was evident visually by a shift in correlation function towards left (Figure 5B). Interestingly at 300mM of arginine concentration, r_H_of the protein was found to be 19.5Å, which was similar to the expected rH of a globular (with well-defined secondary structure) protein of 140 amino acid residues (17.7-19.9Å).

Since the discrete diffusional components method, which was used in the above FCS analyses, is model dependent, correlation functions were further analyzed using MEMFCS. MEM distributions determined at different arginine concentrations showed a large shift towards left indicating a decrease in τD (Figure S5). We found that both methods of analysis (MEMFCS and discrete diffusional components) yield similar values for the extent of arginine-induced compaction. The conformational fluctuation of α-syn between the extended and compact conformers has been suggested before[22, 23]. A measure in an average number of particles (N) with arginine showed no change in N as shown in Figure 5D, indicating no protein aggregation at this solution condition. Aggregation is expected to result in a decrease in N (Figure 5E), as shown before [24].

In contrast to arginine, the addition of glutamate shifted correlation functions towards right (Figure 5F). The data in the presence of glutamate could not be fit to a single component diffusion model (non-random nature of the residual distribution shown in Figure S6A). The data was instead fit to a model (with random residual distribution, FigureS6B), which contained two diffusing components (equation 8). The τD of the faster component (τD1) was fixed to the value of the monomer, while the second component was found to be a slowly diffusing oligomer (τD2). The amplitude of the first component (monomer, a_1_) decreased, while that of the oligomeric component (a_2_) went up as the concentration of glutamate increased (Figure 5G). The value of r_H2_ also increased with glutamate concentration (Figure 5H). The value of N decreased with the increase in glutamate concentration (Figure 5I), supporting glutamate induced oligomerization (Figure 5E). Count per particle analysis was however complicated because of aggregation induced quenching and loss of bright aggregates from the analysis in the form of spikes [24]. At glutamate concentration greater than 200mM, the data could not be fit using two components because of greater heterogeneity of the aggregation process (Figure S6C). The best fit was achieved using a three component model (Figure S6C). Independent MEMFCS analyses (equation 12) showed an increase in the values of τD (Figure S7). The prominent broadening of the MEM profiles in the presence of glutamate (Figure S6), presumably suggested the presence of multiple unresolved diffusing components [24]. We used FTIR spectroscopy to obtain insights into the secondary structure of the glutamate induced oligomers. FTIR data showed the presence of disordered segments in the native condition (Figure S8A), the extent of which decreased significantly as oligomers started forming in the presence of glutamate. FTIR data clearly showed that the oligomers contained significant extent of anti-parallel β-sheet characters. Glutamate-induced oligomers did not show significant ThT binding (Figure S8B).

Since FCS may not be an appropriate method for large oligomers, we used AFM for their direct visualization. We found that relatively small oligomers (with diameters varying between 10nm and 15nm) predominated at glutamate concentration less than 100mM and in the absence of incubation. In contrast, AFM measurements with glutamate treated sample after 3 hours of incubation showed at least two kinds of oligomeric materials (80% population with 20-35 nm in diameters and 2-4 nm in height and 20% populations with 60-85 nm in diameters, 6-10 nm in height) (Figure S8C). Although glutamate induced oligomers contained significant extent of anti-parallel β-sheets, they did not show ThT binding. In the presence of arginine, we did not find any oligomers even after 3 hours of incubation at 37°C. It may be noted that the size of the early oligomers has been found to vary significantly depending on the experimental conditions [25]. For example, the oligomers isolated from the post-mortem brains can be typically divided into small (∼2–5 mers), medium (∼5–15 mers), and large (∼15–150 mers) categories[26, 27]. Single molecule FRET experiments showed small, medium and large oligomers, which were populated before the generation of fibrils [27].

### Arginine and glutamate cancel each other when used in a mixture

We found that the presence of glutamate enhanced fibrillation (the late stage of aggregation) presumably by favoringoligomerization at the early stage. In contrast, arginine inhibited fibrillation by inducing compaction. Subsequently, we wanted to find out if these properties were additive for both early and late stages. As before, we carried out two sets of experiments, the early events using FCS, the late events using ThT fluorescence and live cells microscopy.

Since the diffusion behavior of the protein is different in glutamate and arginine, and hence we used a mean diffusion time approach to analyze FCS data (Supporting Information). In the presence of initially added arginine, the value of r_Hm_ started decreasing as the protein attained a compact structure (Figure 6A). In the presence of glutamate, which was added in the presence of 100mM arginine, the values of r_Hm_ (equation 14) started increasing as a result of glutamate induced oligomerization (Figure 6A). Independent MEM analyses showed identical trend (Figure S8A). The data showed that the addition of glutamate in the mixture can efficiently reverse the effect of arginine. Figure 6B showed the result of the reverse experiment in which the addition of arginine in the mixture countered the effects of glutamate. An arginine to glutamate ratio of 2:1 could be established at which condition these two co-solvents completely canceled each other’s influence. This inference was further supported by plotting Δr_Hm_ (Figure 6C), whose value was approximately zero at the arginine to glutamate ratio of 2:1. We repeated the above experiment using different concentrations of arginine and glutamate keeping the ratio constant. We observed that the value of r_Hm_remained identical as long as the ratio of arginine to glutamate ratio was maintained at 2:1 (Figure 6D). Independent MEMFCS analysis of the correlation functions also showed identical profile distributions (Figure S8B). The 3D contour plot shown in Figure 6E clearly defined a region (represented by cyan) at which the protein behaved like a monomer in aqueous buffer although the solution contained variable concentrations of glutamate and arginine. The residual distributions of FCS fit for different combinations of arginine and glutamate concentrations (keeping the ratio constant) were shown in Figure S6D.

**Figure 6:**
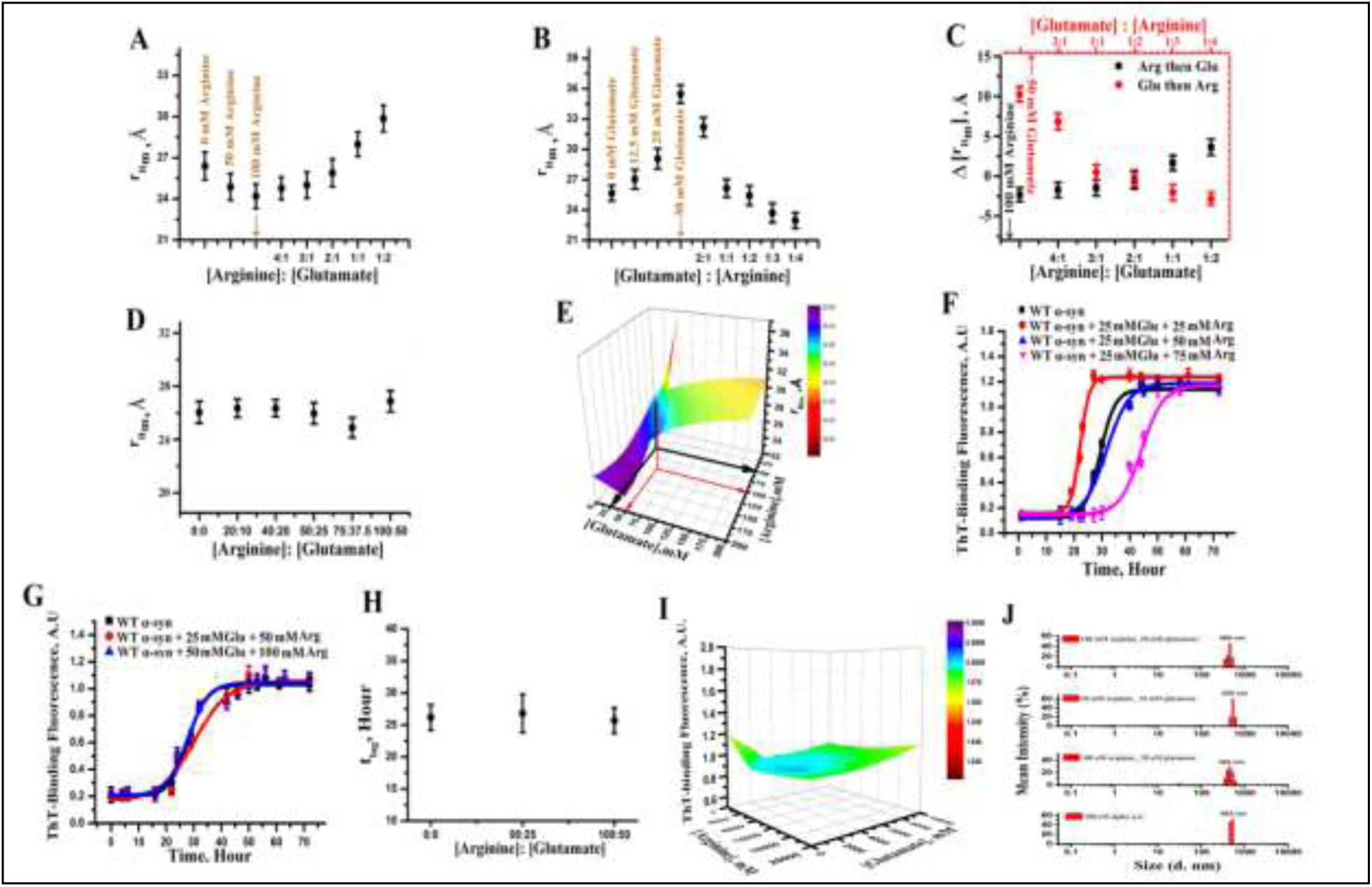
[A] The variations of mean hydrodynamic radius (r_Hm_, Å) with concentrations up to 100mM of arginine, followed by addition of different concentrations of glutamate; [B] the variations in r_Hm_with glutamate concentration up to 50mM, followed by addition of arginine of increasing concentrations; [C] the variation in ΔrHmat different ratio of arginine:glutamate (black) and glutamate:arginine(red). ΔrHm has been calculated by subtracting the value of r_Hm_ at a particular osmolyte (arginine or glutamate) concentration from that observed in the absence of osmolyte; [D] r_Hm_ at different concentrations of arginine and glutamate mixture keeping the ratio constant at 2:1. [E] 3D contour plot using the values of r_Hm_ of both experiments of [A] and [B]; The color cyan indicates the values of r_H_ at the concentration ratios of 2:1 (corresponding to 50mM, 25mM of arginine, glutamate shown by the black arrow; and 100mM, 50mM arginine, glutamate shown by the red arrow) (red) are similar to the native form of protein in the absence of any co-solvents. Color scale bar was shown beside the plot. [F] Amyloidosis kinetics of α-syn at different ratios; 1:1 (red), 2:1 (blue) and 3:1 (pink)] of arginine and glutamate are shown. The data of WT α-syn in aqueous buffer in the absence of any osmolytes(black) is shown as control data set; [G] Amyloidosis kinetics of α-syn with different concentrations of arginine and glutamate (50:25 as red and 100:50 as blue) keeping the ratio constant at 2:1; [H] the variations of t_lag_ of amyloid formation for these mixtures with fixed ratio of 2:1. The data clearly show that arginine inhibits and glutamate facilitates amyloidosis kinetics and the arginine to glutamate ratio of 2:1 cancels each other’s influence completely; Data shown are mean±standard error. Error bars of the FCS experiments were calculated after repeating same experiments independently for five times. [I] 3D contour plot shows variation of ThT-binding fluorescence at saturated condition with different concentrations of arginine and glutamate keeping 2:1 ratio constant. [J] DLS measurement of size of WT α-syn at saturated condition with 2:1 ratio of arginine and glutamate.

The validity of the arginine to glutamate ratio was also established for the late stage using ThT binding assay (Figure 6F). The amyloid formation kinetics remained superimposable for several combinations of arginine and glutamate concentrations, but with the identical ratio of 2:1 (Figure 6G, 6H). Figure 6I shows 3D contour plot of ThT fluorescence intensity data at the saturated conditions of the aggregation kinetics obtained for different concentrations of arginine and glutamate (keeping the ratio constant at 2:1). For plotting the ThT fluorescence intensity at the saturating concentrations, we varied the concentrations from 500μM and 200mM, and the ratio was maintained in the entire range. DLS measurements with different concentrations of arginine and glutamate at 2:1 ratio show similar trends (Figure 6J). The live cells data are shown in Figure 7, which show that after treatment with arginine in the previously glutamate-treated cells, the extent of aggregation was almost similar to that of TC-AS expressing cells without any treatment. In the presence of 2:1 arginine-glutamate mixture, all the three parameters (the diameter of an individual aggregated spot, the maximum intensity of an individual spot, and the integrated area of the inclusion) were calculated to be similar to those observed without any amino acid treatment (Table 1).

**Figure 7:**
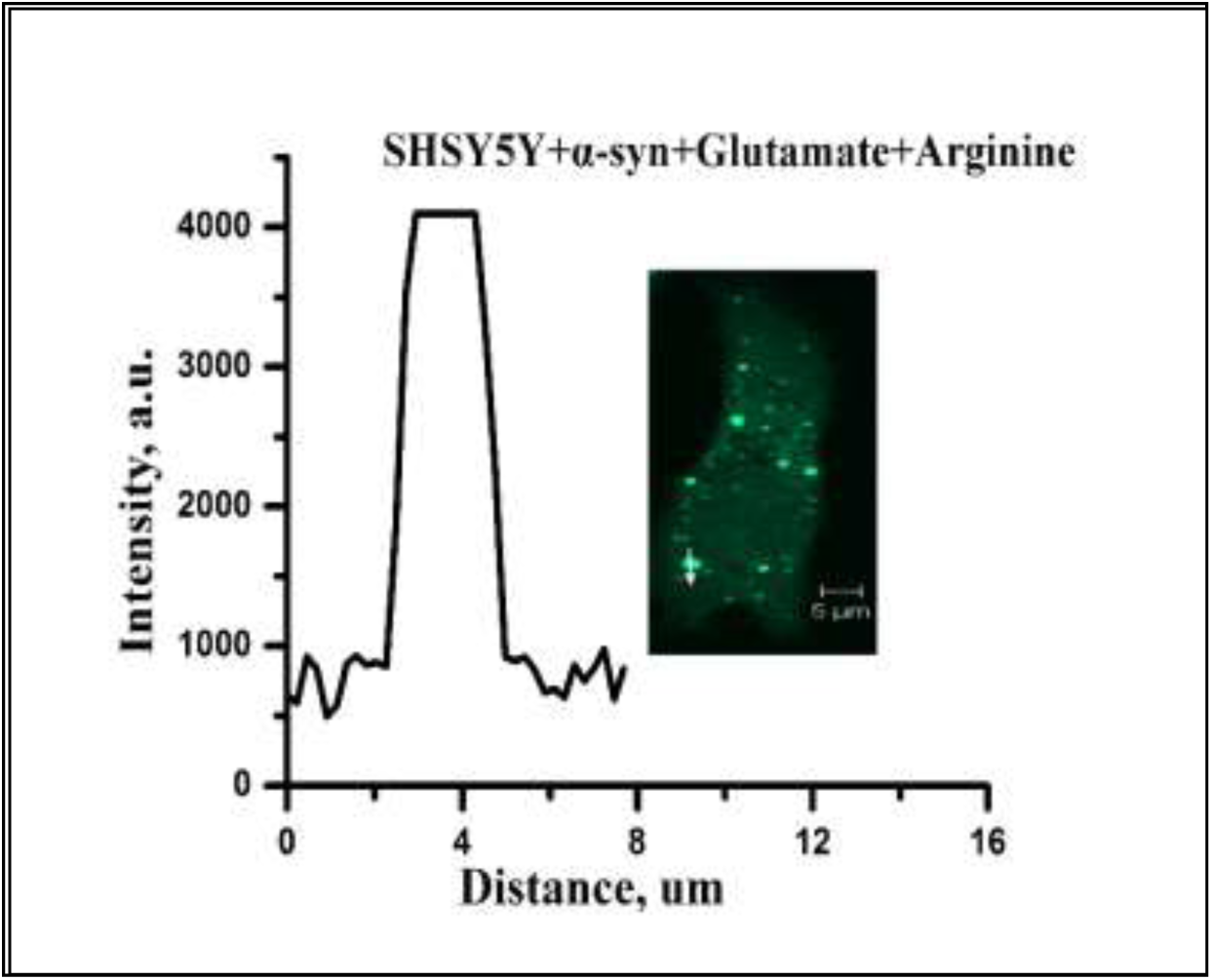
SH-SY5Y cells containing TC-AS aggregates in the presence of both arginine and glutamate at 2:1 ratio. Scale bars, 5μm. The imaging was performed using n=200 TC-AS overexpressing cells, one of them is shown above as a representative.

### Arginine binds to the protein, while glutamate excludes itself from the protein surface

To investigate further the binding between the protein and arginine, we used mass spectrometry and isothermal calorimetry (ITC). ITC data (Figure 8A) clearly suggested a two steps sequential binding between α-syn and arginine, which was found to be enthalpically unfavorable, but entropically driven ((Δ*S*_*1*_ in this case is +51.1 cal/mol/K and Δ*S*_*2*_ = 699 cal/mol/K). The displacement of ordered water molecules by arginine is expected to be enthalpically unfavorable which would involve breaking of multiple hydrogen bonds. Since arginine has been reported to form well-arranged cluster structure through π-stacking interaction through its guanidium moiety [28], the internal energy is larger for this kind of already existing well-packed structures. This is because of the relatively strong cohesive forces that may exist in such cluster structure [29]. Relatively higher endothermic ΔH suggested the probability of the replacement of strong arginine-arginine molecular interaction with protein-arginine interaction, especially in the well-ordered cluster system and the process is favorable (ΔG= −5.6 Kcal/mol) (equation 16). ITC experiments in the presence of glutamate did not show any significant heat flow suggesting very poor or no binding (Figure 9B). The detailed understanding of the binding interactions between α-syn and arginine is being studied in our laboratory. The presence of arginine cluster could be established further by mass spectrometry. The data in the absence of protein clearly suggested the presence of multimeric arginine species (Figure 9C), whose intensity decreased once α-syn was added (Figure 9D).

**Figure 8:**
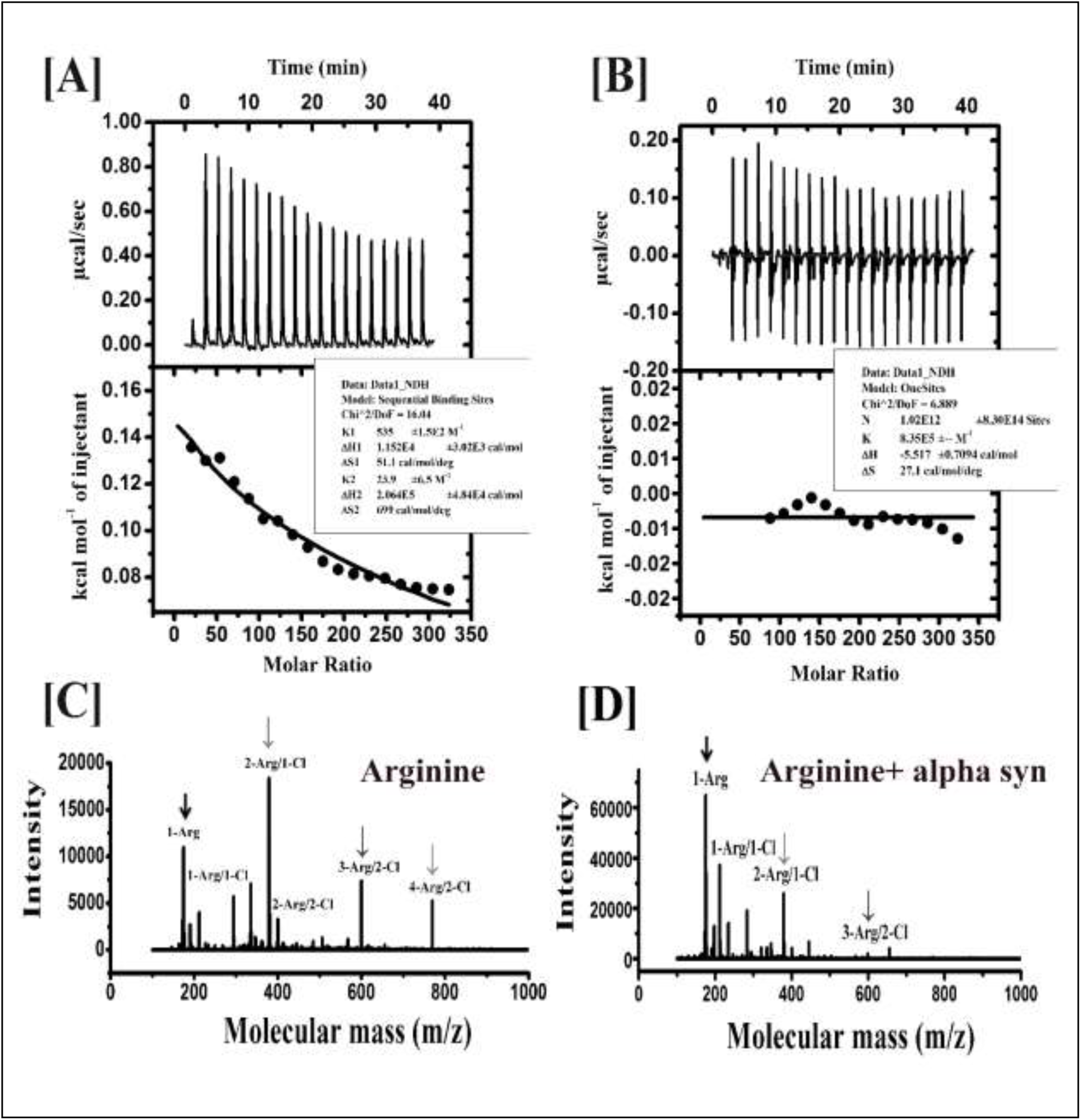
[A] ITC thermogram and binding graph of α-syn with arginine. Δ*G* of binding has been determined to be-4 kcal/mol and −1.6 kcal/mol. The total value of Δ*G* (-5.6 kcal/mol) provides a quantitative measurement of binding strength. Δ*S*_*1*_ in this case is +51.1 cal/mol/K and Δ*S*_*2*_ = 699 cal/mol/K, which are positive. [B] the titration profile of the protein with glutamate (same procedure as with arginine), indicating random heat flow with the increase of the number of injections. [C] Stable arginine-arginine interaction has been evident from its high molecular weight multimer (cluster) formation in the solution phase, which are indicated by arrows. [D] the replacement of arginine-arginine interaction with protein-arginine interaction is evident from the decrease in intensity of the large clusters (shown by arrows) from MALDI-TOF.

Figure 9 shows a mechanistic outline of the effect of arginine and glutamate on α-synfolding-aggregation landscape, which we developed from the presented data obtained in the aqueous buffer. The role of a small molecule depends on many factors including a competition between direct binding and preferential exclusion. An osmolyte like glutamate may not bind to a protein; rather they prefer to remain excluded. The aggregation facilitation property of glutamate presumably comes from preferential exclusion from the protein surface, because of which glutamate would favor conformational states with less surface area facilitating aggregation (as aggregation reduces overall surface area). The use of aspartate (which has the same charge) did not seem to have any major effect supporting the hypothesis that there was no significant binding. However, it is difficult to explain the difference in behaviour between glutamate and aspartate-primarily because of the fact that the osmophobic property of aspartate is not well studied. In contrast, the presented ITC results supported arginine binding to the protein. Arginine has been found to weakly bind to protein molecules through aromatic residues using cation-pi interactions [30]. The importance of the guanidinium moiety, which is present in arginine, has been noted by the fact that lysine (with similar charge and no guanidinium moiety) did not inhibit α-syn aggregation (Figure 4A). In contrast, guanidinium hydrochloride was found to inhibit aggregation (Figure 4A). The apparent difference in the dose dependence of guanidinium hydrochloride and arginine is not explained completely at present, but it should be noted that guanidinium hydrochloride is a strong denaturant while arginine is not. The involvement of supramolecular assembly of arginine towards protein binding has been suggested [31]. Similar weak interaction may play important roles in the use of small polyphenolic compounds, e.g. curcumin, which has shown the prevention of α-syn aggregation [32]. The presented results also suggested strongly that the effect of arginine was not due to the presence of salt, as the addition of salt facilitated aggregation. The salt behaviour observed in the present study matches with already published data on α-syn[33].

**Figure 9:**
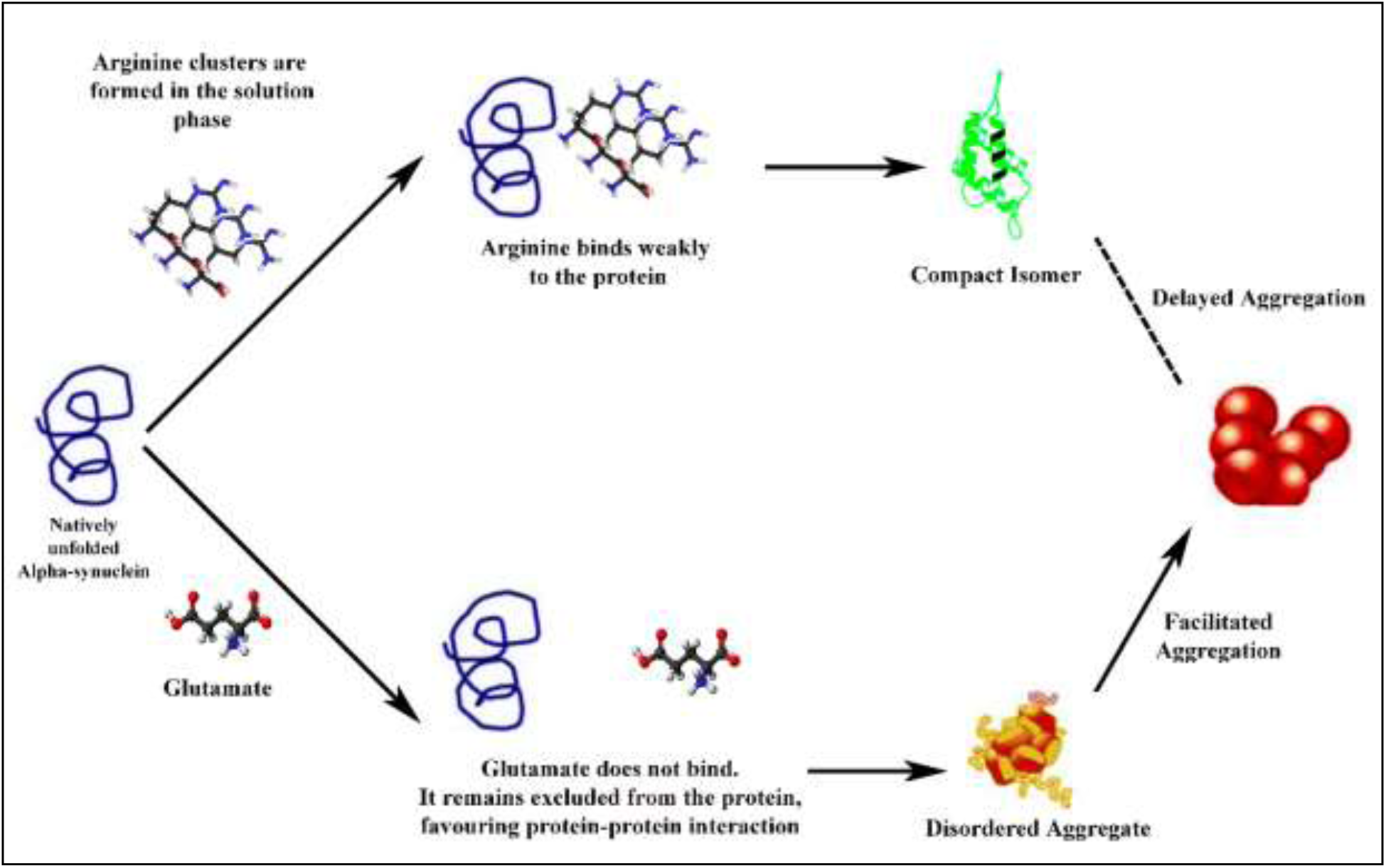
Mechanistic outline of the effect of arginine and glutamate on α-syn aggregation landscape

The major objective of this study was to find out how three components of α-syn aggregation modulate each other. When fibrillation was inhibited by arginine, we found that the protein preferred to populate the compact isomer in the early stage (Figure 9). In contrast, the facilitation of the late stage aggregation by glutamate favored early oligomerization (Figure 9). The use of small molecules, particularly osmolytes, has been found to modulate aggregation in several neurodegenerative disease systems. For example, TMAO and glycerol are shown to prevent the conversion of PrP^C^ into PrP^Sc^in mouse neuro-blastoma cells[34]. Glycerol is found effective in reducing the lysosomal accumulation of toxic prion proteins[35]. 7.5% solution of DMSO, when administered by oral route, has been shown to delay the accumulation of PrP^Sc^ in hamsters intra-cranially inoculated with prions[36]. Oral administration of 2% trehalose solution has been shown to minimize aggregation of protein Huntington to improve motor dysfunction and extend lifespan in a transgenic mouse model of Huntington Diseases[37].

The present paper highlights an important advantage of a small molecule based approach in which a reverse phenotype can easily be generated by using an opposing small molecule.The effect of mixed co-solvents on protein conformation and stability has been investigated before. A mixture of urea and sarcosine has been used to study unfolding co-operativity (m-values), and the results clearly show that each osmolyte can act independently[38]. Glutamate and TMAO have been found to counteract the destabilizing effect of urea [39, 40]. The expansion and contraction of α-syn were found to cancel each other at a 2:1 ratio of TMAO and urea. While the additive property of small molecules has been well studied to decipher multiple aspects protein conformation and stability, the present study suggests that this addictiveness can also be extended to protein aggregation.

Although the studies of protein folding and aggregation are traditionally carried out in dilute aqueous solution, the situation is rapidly changing. The advancement of experimental techniques, particularly NMR [41] and fluorescence methods[42] now result in efficient probing of the complexity of the cellular systems possible. Although the present data with small molecules show interesting correlation between the observations in aqueous buffer and inside live cells, a complete understanding of this would require detailed and extensive investigation, which is currently the present focus of our laboratory.

## EXPERIMENTAL PROCEDURES

### Materials

Isopropyl β-D-galactopyranoside (IPTG) and Tris salt were purchased from Biotecx Laboratories (Houston, TX) and J.T. Baker (Center Valley, PA) respectively. Phenylmethanesulfonylfluoride (PMSF), ammonium sulfate, and sodium chloride were obtained from Sigma-Aldrich (St. Louis, MO). Arginine hydrochloride and Glutamate sodium salt were obtained from Sigma-Aldrich (St. Louis, MO). FBS, DMEM, Opti-MEM were obtained from Life Technologies (Carlsbad, CA). All experiments were carried out using 20mM sodium phosphate buffer at pH 7.4 unless otherwise mentioned. Special care was taken to maintain the pH of the solutions, when small molecules (glutamate, aspirate, lysine or arginine) were used.

The expression, purification and fluorophore labeling (where applicable) of WT and mutant proteins have been carried out using published procedure [20]. TC-AS construct was a generous gift from Prof. Donna Arndt-Jovin (Max Planck Institute for Biophysical Chemistry, Laboratory of Cellular Dynamics).

### DNA Constructs for the Live Cells Studies

For these studies, we used a small ∼ 1.5kDa tetra-cysteine tagged α-syn (TC-AS) construct, which was a generous gift from Prof. Donna Arndt-Jovin (Max Planck Institute for Biophysical Chemistry, Laboratory of Cellular Dynamics). A 12mer peptide was fused with the C-terminal of the protein and cloned in pcDNA3.0[16].

### Cell Culture, Transfection and Treatment

We have used a cervical cancer cell line (HeLa) and a neuroblastoma cell line (SH-SY5Y) for these measurements. Both HeLa and SH-SY5Y cell lines were maintained in DMEM supplemented with 10% heat-inactivated fetal bovine serum (FBS), 110 mg/L sodium pyruvate, 4mM l-glutamine, 100units/ml penicillin and 100μg/ml streptomycin in humidified air containing 5% CO_2_ at 37 ° C.

Cells were transiently transfected with 2.5μg wild type TC-AS construct using Lipofectamine LTX and Plus reagent (Invitrogen, Carlsbad, CA) as described by the manufacturer. 24 hours before the transfection, cells were seeded on 35mm poly-D-lysine coated plate (MatTek Corporation, Ashland, MA) and allowed to grow to ∼60% confluent. For the treatments, TC-AS transfected cells were incubated with 500mM of co-solvents (arginine or glutamate) supplemented in growth media for 20 hrs. Before imaging, cells were stained with TC-Flash (Life Technologies, Carlsbad, CA) as per manufacturer’s protocol. Nuclear staining was performed using Hoechst 33342 in PBS buffer to identify the living Hela and SH-SY5Y cells.

### Thioflavin T (ThT) Binding Fluorescence Assay

Amyloidosis kinetics of α-syn was measured using the help of ThT fluorescence assay using a PTI fluorimeter (Photon Technology International, NJ). 500 μl of 150μM α-syn was agitated 180rpm at 37°C. At different time points of incubation, fluorescence intensity was measured for the samples containing 1.5μM α-syn and 20μM of ThT in 20mM sodium phosphate buffer pH 7.4. Slits for this experiment were 5 nm for both excitation and emission, while the integration time was 0.3s. Typical excitation and emission wavelengths were 450 nm and 485 nm respectively. The path length of the quartz cuvette was 1 cm. Aggregation kinetics data were fit to the following equation[43]:

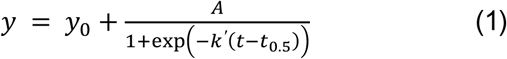

Where y_0_ was the signal base line at the lag phase; A was the total increase in fluorescence signal between the lag and stationary phase;k’ is the growth rate constant and t_0.5_ was its mid-point of the log phase. From this lag time is calculated by the following equation.

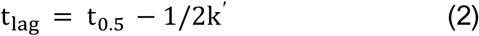

We calculated t_lag_of aggregation in the presence of different co-solvents, and this value was used as a measure of the aggregation propensity.

### Dynamic Light Scattering (DLS)

The Malvern Zetasizer Nano ZS DLS system (Malvern Instruments Ltd., UK) was used to perform all DLS measurements. This system is equipped with a 633 nm He–Ne laser and an APD detector configured to collect backscattered light at 173°. The measurement sample was allowed to equilibrate for 120 s at 20°C prior to the measurements. Fifteen separate runs, each of 10s duration were averaged for each DLS measurement. DLS data were analyzed using Malvern Zetasizer 6.12 software. The reported mean particle hydrodynamic radii (R_H_) were calculated using the intensity based particle size distributions.

### Atomic Force Microscopy (AFM)

AFM was performed using a Pico Plus 5500 AFM (Agilent Technologies, USA) with a piezoscanner having a maximum range of 9 μm. 10 μL of the samples were deposited on a freshly cleaved muscovite ruby mica sheet (ASTM V1grade ruby mica from MICAFAB, Chennai). After 30 minutes the sample was dried with a vacuum drier. The cantilever resonance frequency was 150–300 kHz. The images (256 pixels × 256 pixels) were captured using a scan size of between 0.5 and 8 μm at a scan speed of 0.5 lines/s. The length, height, and width of protein fibrils were measured manually using PicoView1.10 software (Agilent Technologies, USA).

### Confocal Microscopy

These experiments were carried out using a Zeiss LSM510 Meta confocal microscope equipped with C-Apochromat 40X (NA=1.20, water immersion) objective and confocal images were acquired with 512X512 pixel (pinhole aperture ∼ 1 airy units). Flash-EDT_2_ dye was excited using an argon laser at 488nm. Hoechst 33342, for nuclear counterstaining was excited with a 405nm diode laser. A 563nm laser was used to excite Lysotracker red. All live cell imaging was performed in 20mM HEPES-based medium.

For the FRAP measurements, the scanning laser power was set to 5%, whereas the bleaching at the region of interest (ROI, ∼2μm in diameter) was carried out using 100% of the laser power. Pre-bleaching was carried out by taking 10 images, which was followed by photo-bleaching with 20 iterations at 100% laser power. Eighty (80) post-bleaching images were acquired.

### FRAP Analysis

Fluorescence recovery curves were constructed from the total intensity fluorescence values at ROI for each frame of the temporal sequence, corrected for the background and laser scanning bleaching, and normalized to the immediate post-bleached value (F_0_). The mobile fraction associated to each ROI (F_M_) was estimated using the following Equation from the fluorescence intensity values before photo-bleaching (F_i_), immediately after photo-bleaching (F_0_), and at the end of the experiment (F_∞_) [17]:

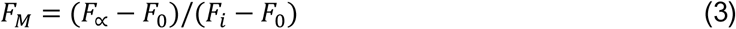

The recovery data was fit to the following equation[44]:

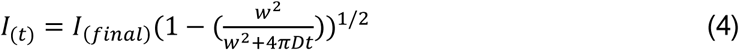

Where, I_(t)_ is the observed intensity as a function of time; I_(final)_ is the final intensity, which was reached after recovery; w^2^ is the ROI area; and D is the value of diffusion coefficient. The values of D were determined by fitting experimental FRAP data to equation (4).

### Additional Data and Image Analyses

Confocal images were analyzed and the diameter and areas of fluorescent punctates were calculated using ImageJ software (NIH, MD). Further image analysis and graphical representations were performed using OriginPro 8.5 (OriginLab Corp., MA)

### Labeling of α-syn with Alexa488Maleimide

α-syn does not contain any cysteine residue. We introduced a cysteine mutant (G132C) at the c-terminal end of the protein. G132C mutant behaves exactly like wild type [20]. This cysteine mutant was labeled with Alexa488Maleimide (Alexa488) using a published procedure [45]. We removed the excess dye using extensive dialysis, which was then followed by column chromatography using a Sephadex G20 column equilibrated with 20 mM sodium phosphate buffer (pH 7.4).

### Fluorescence Correlation Spectroscopy (FCS)

FCS experiments were carried out using LSM 510 Meta (Carl Zeiss, Germany) with 40X water immersion objective. The labeled protein samples were excited using an argon laser at 488 nm. Fluorescence signals were separated from the excited line by dichroic filter and then collected using two avalanche photodiodes (APD). Correlation functions were calculated from the photocurrent detected by these APDs. Refractive index and viscosity related effects of the osmolyte solutions were minimized and normalized using previous methods[46].

In an FCS experiment, fluorescence intensity fluctuations are recorded as the fluorescently labeled molecules diffuse in and out of the confocal volume. The fluctuations in fluorescence intensity consisting of information on the average number of molecules (N), the residence time and other photo-physical properties can be analyzed using a suitable model.

The normalized form of the autocorrelation function of fluorescence intensity I(t) at time t is represented by:

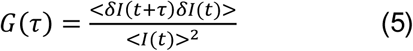

In this expression, <*I* (*t*)> is the average of fluorescence signal over time, and *δI*(*t*) is the signal fluctuation at time *t* minus the average:

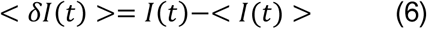

For a simple solution containing a single diffusing species, the correlation function can be represented as follows:

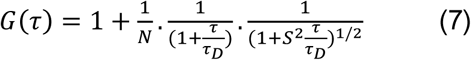

Where τ_D_ is the characteristic diffusion time, N is the average number of particles in the observation volume, and S is the structural parameter.

For a more complex system containing multiple diffusing species (excluding the triplet state contributions), the correlation function can be defined by:

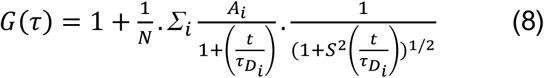

Where τDi is the diffusion time of the i^th^ diffusing species present in the solution and Ai is its relative amplitude. It is important to note that

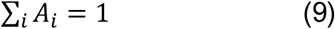

Where τ_Di_ is the diffusion time of the i^th^ component and A_i_ is its amplitude in the correlation function.

The value of τD obtained by fitting the correlation function is related to the diffusion coefficient (D) by the following equation:

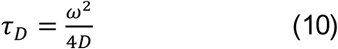

The value of ω, which defines the size of the observation volume, was calculated using FCS measurements with Rhodamine 6G, whose value of D has been well established[47]. The value of the hydrodynamic radius (r_H_) can be obtained from D using the Stokes’ Einstein formalism (Equation11).

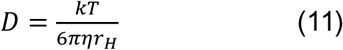

Where η is the viscosity and k is the Boltzmann constant. It may be noted that Equation (11) assumes that the molecules under investigation are spherical.

The presence of glutamate and arginine results in an increase in the refractive index of the measurement solutions. We minimized the effect of refractive index and viscosity on the correlation functions using previously published method [46]. First, we optimized the FCS data by suitably changing the collar settings and the height of the objective. Second, the protein data were normalized using the values of □ _D_ obtained with the free dye (Alexa488) measured under identical conditions.

### Maximum Entropy Method (MEM) Analysis

Correlation functions were analyzed further using MEMFCS algorithm, which has been recently developed to produce a model free understanding of FCS data [48]. MEMFCS determines αidistribution of a heterogeneous system in an unbiased way on the basis of maximum entropy S and minimum χ2. Maximum entropy (S) is defined by-

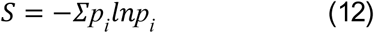

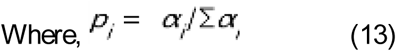

### The determination of mean diffusion time (t_Dm_) and mean hydrodynamic radii (r_Hm_) in the presence of both small molecules

The diffusion behavior of the protein in the presence of both small molecules was studied using FCS. First, the diffusion behavior of Alexa488Syn was studied by FCS in the presence of arginine concentration varying between 0 and 100mM. As described in the text, FCS data with arginine were analyzed using a single component model. In the presence of 100mM arginine, glutamate was added and its concentration was varied between 25 and 200mM.As glutamate concentration increased, FCS analyses started becoming complex with the fits deviating from the single component model. To show the results obtained using two different models (one component vs. two components in the presence of arginine and glutamate respectively), we used a mean t_D_ approach.

We defined the mean diffusion time (t_Dm_) as:

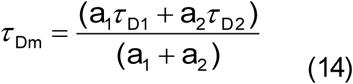

The values of mean diffusion coefficient (Dm), and mean hydrodynamic radii (r_Hm_) were calculated from the values of τDm using equations (10) and (11).

### FT-IR Spectroscopy

FT-IR spectra of α-syn at different conditions were acquired by using Bruker Tensor 27 spectrometer. Buffer baseline was subtracted before taking each spectrum. The assignment of peaks at amide I (1600-1700cm^-1^) was done using previously described spectral components associated with different secondary structures [49].

### Isothermal Titration Calorimetry (ITC)

VP-ITC micro-calorimeter was used in high feedback mode to measure the heat flow resulting from the binding of α-syn with both amino acids used here; arginine and glutamate. α-syn was dissolved in the 20 mM sodium phosphate buffer and pH of arginine and glutamate solution was maintained in 7.4. Sample cell with 25 μM α-syn was titrated against 40 mM of amino acids (arginine or glutamate) in a syringe with 300 rpm stirring speed. The cell temperature was kept at 298 K and 10 μcal/s of reference power as applied to maintain a flat baseline. A total of 19 injections of 40 mM arginine/glutamate was performed, where the initial injection was of 0.5 μl and the final was 2.0 μl, while the remaining 17 injections were 2 μl each. Each injection was over a period of 0.8 sec with a spacing of 70 sec and filter time of 5 sec.

The heat of dilution, *h*0, was determined in control experiments by injecting the corresponding amino acid dispersion (or arginine/glutamate solution) into the buffer solution. The heats of dilution (not shown) were subtracted from the heats determined in the corresponding protein-amino acid binding experiments. Thus, the quantitative evaluation of the experimental data was based on the relationship, Δ*H* = *h*i – *h*0. The overall binding enthalpy and the binding isotherm were determined using standard procedures [50]

Results obtained were plotted using MicroCal Origin 7.0 software. In order to determine the thermodynamic parameters (i.e., association constant K_A_ and enthalpy of reaction ΔH), the single-site binding model was used. Other thermodynamic parameters such as the free energy of binding (ΔG) and entropy (ΔS) were calculated using equations (15) and (16), respectively:

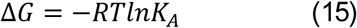

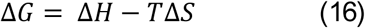

### MALDI-TOF Analysis

For MALDI-TOF analysis we used 100 mM of arginine solution (pH 7.4) with varying concentration of α-syn from 10μM to 40 μM. 0.5 ml was spotted on a-cyano-4-hydroxy-cinnamic acid matrix. Analysis of the sample was performed using MALDI-TOF by an Applied Biosystems Q10 4800 MALDI TOF/TOF™ analyzer using 4000 series explorer software for acquisition and GPS explorer software, version 3.6 for analysis. 4700 calmix (Applied Biosystems) were used to calibrate the instrument under reflector mode. The error limit is 50ppm.

### MTT Assay

A colorimetric MTT metabolic activity assay was used to determine cell viability. HeLa and SH-SY5Y cells (1 × 10^4^cells/well) were cultured in a 96-well plate at 37 °C. After treatment, MTT reagent (Invitrogen, Carlsbad, CA) was added and incubated for 4hrs at 37°C. The resultant formazan crystals were dissolved in dimethyl sulfoxide and the absorbance was measured at 540 nm using a microplate reader

## ACKNOWLEDGMENT

AK and SG contributed equally to this work. We are grateful to Prof. Donna Arndt-Jovin (Max Planck Institute for Biophysical Chemistry, Laboratory of Cellular Dynamics) for providing us the TC-AS construct. We thank Prof. Sudipta Maiti for providing us the MEMFCS algorithm. The funding from the Department of Science and Technology (EMR/2016/000310) has been acknowledged. AK and SG thank UGC for awarding fellowships. We thank the Director, CSIR-IICB for his encouragements. We thank Mr. Sandip Chakraborty (CSIR-IICB) for the MALDI-TOF/TOF measurements. We thankfully acknowledge DBT-CU IPLS for ITC experiments.

### Conflict of Interest Statement

No author has an actual or perceived conflict of interest with the contents of this article.

### Author Contributions Statement

AK and SG carried out the experiments and analyzed the data. KC conceived the idea, coordinated the study and wrote the paper. All authors approved the final version of the manuscript. AK and SG contributed equally.

